# Evolutionary analysis of p38 stress-activated kinases in unicellular relatives of animals suggests an ancestral function in osmotic stress

**DOI:** 10.1101/2022.04.24.489300

**Authors:** Victoria Shabardina, Pedro Romero Charria, Gonzalo Bercedo Saborido, Ester Diaz-Mora, Ana Cuenda, Iñaki Ruiz-Trillo, Juan Jose Sanz-Ezquerro

## Abstract

p38 kinases are key elements of the cellular stress response in animals. They mediate the cell response to a multitude of stress stimuli, from osmotic shock to inflammation and oncogenes. However, it is unknown how such diversity of function in stress evolved in this kinase subfamily. Here, we show that the p38 kinase was already present in a common ancestor of animals and fungi. Later, in animals, it diversified into three JNK kinases and four p38 kinases. Moreover, we identified a fifth p38 paralog in fishes and amphibians. Our analysis shows that each p38 paralog has specific amino acid substitutions around the hinge point, a region between the N-terminal and C-terminal protein domains. We showed that this region can be used to distinguish between individual paralogs and predict their specific. Finally, we showed that the response to hyperosmotic stress in *Capsaspora owczarzaki*, a close unicellular relative of animals, follows a typical for the p38 kinases pattern of phosphorylation-dephosphorylation. At the same time, *Capsaspora*’s cells upregulate the expression of GPD1 protein resembling an osmotic stress response in yeasts. Overall, our results show that the ancestral p38 stress pathway originated in the root of opisthokonts, most likely as a cell’s reaction to salinity change in the environment. In animals, the pathway became more complex and incorporated more stimuli and downstream targets due to the p38 sequence evolution in the docking and substrate binding sites around the hinge region. Overall, this study improves our understanding of the p38 evolution and opens new perspectives for the p38 research.

## Introduction

Stress response in multicellular organisms is a complex process essential for cell survival. The p38 family of stress activated kinases is a major player in this process in animals and is considered to be animal-specific [1]. p38 kinases belong to the larger group of mitogen-activated protein kinases (MAPKs). In animals, they are activated in response to almost all stress factors, from osmotic shock and UV radiation to cytokine-mediated inflammation and oncogene activation [2]. They also regulate apoptosis, cell differentiation, proliferation, and migration [3]. Not surprisingly, dysregulation of p38 mediated pathways can lead to a number of pathologies, including inflammatory, cardiovascular and neurodegenerative diseases, and cancer.

Vertebrates have several p38 paralogs with seemingly functional redundancy, which is a typical example of paralogs evolution in vertebrates. Thus, most invertebrates have one gene coding for p38 kinase [4], [5] with some exceptions [6], [7], while vertebrates are known to have four paralogs: p38α (MAPK14), p38β (MAPK11), p38γ (MAPK12), and p38δ (MAPK13). The vertebrate paralogs share high sequence similarity, and their functional redundancy was suggested in several studies [8], [9]. However, there is also evidence advocating individuality and specific biological roles of each paralog. For example, they may differ in substrate specificity and specificity to their activating kinases (MAPK kinases or MAPKKs). Different p38 paralogs can also exhibit different sensitivity to kinase inhibitors and tissue expression [10]–[15]. Therefore, it is important to understand the role of each p38 in the cell’s stress response. However, still little is known about the molecular mechanisms behind the paralogs’ individuality.

The four p38 vertebrate paralogs originated by a tandem duplication that led to the presence of p38β and p38γ genes. This latter was followed by segmental duplication that resulted in two more paralogs, p38α and p38δ [6]. Two other closely related groups of kinases, also known for their role in the stress response, are the c-Jun N-terminal kinase (JNK) in animals and Hog1 in fungi. p38, JNK, and Hog1 kinases share a similar domain structure and are thought to have originated from the same ancestral protein [16], [17]. The most conserved parts of these proteins correspond to 1) an ATP binding site, 2) the docking site and the common docking (CD) motif, 3) an activating loop, and 4) a substrate-binding site [18]. The docking site and the activating loop determine the specificity of a MAPK to its substrates and activators. The activating loop also contains a dual phosphorylation motif that is highly conserved in p38, Hog1 (TGY) and JNK (TPY). The dual phosphorylation of the threonine and the tyrosine by MAPKK is characteristic of the stress-activated kinases and is essential for their enzymatic activity [13], [19], [20]. Additionally, there are other motifs that are specific for certain kinases. For example, p38γ contains a PDZ domain-binding sequence at its C-terminal [21]. Less studied are the lipid binding sites which have been described, for example, in p38α. They are thought to be involved in binding specific lipid molecules and modulating the kinase’s activity [22].

Despite its importance, the origin and the evolutionary history of the p38 protein family remain unclear. Existing phylogenetic studies of MAPKs are limited to a narrow set of species, rarely outside of the vertebrate group. Moreover, the prevalence of p38α studies over studies that include the other p38 paralogs further limits our knowledge of the overall network of interactions between different players in the p38-mediated stress response. To fill these gaps, we performed a taxon-rich bioinformatics survey, followed by comprehensive phylogenetic and sequence evolution analyses of the p38 kinase family. Our data show that p38 stress signaling originated before Metazoa. In particular, we identified p38/Hog1 homologs in the genomes of several unicellular relatives of animals, including Choanoflagellata, Filasterea, and Teretosporea. We further characterized expression and phosphorylation patterns of the p38 homolog in the filasterean *Capsaspora owczarzaki* under osmotic stress conditions. Our work shows that p38-mediated stress response in animals might have originated from a relatively simple mechanism activated in the osmotic stress and later, in animals, highly diversified to react to multitude of stimuli. We hypothesize that this diversification likely happened due to the evolution within a specific “hotspot” in the protein’s sequence that we here describe. This work improves our understanding of the molecular basis behind the complex network of p38 signaling and the role of paralog individuality in it.

## Results

### Stress kinases in unicellular relatives of animals

To elucidate the origin of p38, we first performed a bioinformatic survey across all major eukaryotic lineages and then inferred a phylogenetic tree with the protein sequences (Table S1, Supplementary). Our data indicate that p38 and JNK are present in most Metazoa, including the early-branching phyla Porifera, Placozoa, and Cnidaria. Moreover, we also retrieved homologs in several unicellular relatives of animals, namely Choanoflagellata, Filasterea, and Teretosporea (Ichthyosporea + Corallochytrea) (Fig. 1). Interestingly, these lineages are the only non-metazoan eukaryotes whose genomes encode p38 homologs. The corresponding sequences branch in an intermediate position between the animal p38s and JNKs and the fungal Hog1 kinase. Interestingly, the genomes of choanoflagellates encode two p38-like paralogs (named here as choano-1 and choano-2) that makes them the only lineage outside Metazoa with two p38-like proteins. Moreover, choano-1 seems more related to Hog1, while choano-2 appears in a position of a sister group to metazoan p38s. The homolog from *Parvularia atlantis*, a representative of a recently described nucleariids lineage [23], clusters within the fungal Hog1 group. Based on this tree, we hypothesize that the last common ancestor of Opisthokonta (Holozoa and Holomycota) possessed an ancestral protein Hog1/p38 and that was duplicated into the Hog1 gene (in Nucleariida and Fungi) and an ancestral p38-like protein in the root of Holozoa (i.e., the clade comprising animals, choanoflagellates, filastereans, and teretosporeans). The latter evolved into p38 and JNK kinases within the animal clade.

**Figure 1.**
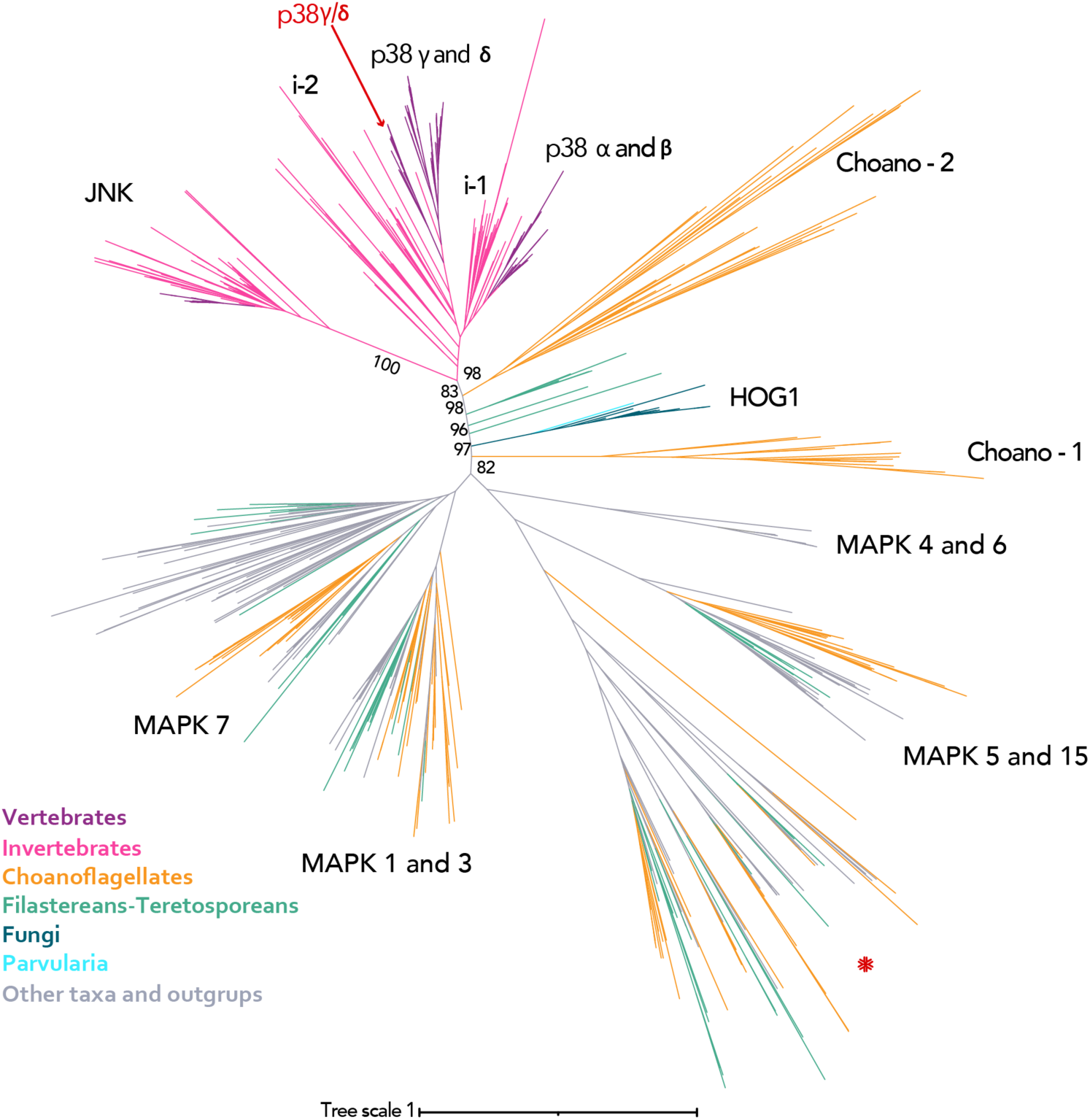
Maximum likelihood phylogenetic tree of the p38 subfamily. Nodal support values (ultrafast bootstrap, 1000 iterations) are shown at the most interesting nodes. The clade named i-1 includes sequences from invertebrate animals such as Arthropoda, Lophotrochozoa, Tunicata, *Trichoplax adhaerens, Intoshia linei, Saccoglossus kowalevskii*. The clade i-2 includes proteins from Arthropoda, Nematoda, Tardigrada, Platyhelminthes, Porifera, and Cnidaria. A new p38 paralog p38γ/δ is highlighted in red. (*) A cluster containing sequences from choanoflagellates, filastereans, teretosporeans, ancyromonads, and the CRuMs supergroup. The identification of this group is out of scope of this study.

We also retrieved several MAPKs from unicellular relatives of animals (orange and green branches in Fig. 1). In particular, choanoflagellates, filastereans, and teretosporeans have MAPKs from all major groups (Table S2, Supplementary) except MAPK4 and MAPK6. The choanoflagellates seem to have the highest content and diversity of MAPKs among unicellular relatives of animals.

Despite the observation that many invertebrates possess only one gene coding for the stress-induced MAPK, there are several cases of lineage-specific duplication. The two well-known examples are *Drosophila melanogaster* and *Caenorhabditis elegans*. The genome of *D. melanogaster* encodes for three p38-like paralogs. One of them, p38c, is characterized by a very long branch on the tree, reflecting a specific role of p38c kinase in fly physiology. Indeed, p38c is known to have a dual functionality: it participates in the stress response typical to p38 group and has a novel function in lipid metabolism [24]. PMK-3 from *C. elegans* is another paralog with a long branch, and it was shown to be a part of a novel mechanism of survival [25], [26]. In this study, we found five more examples of invertebrates with a highly diverged second p38 paralog (Table 1).

**Table 1.**
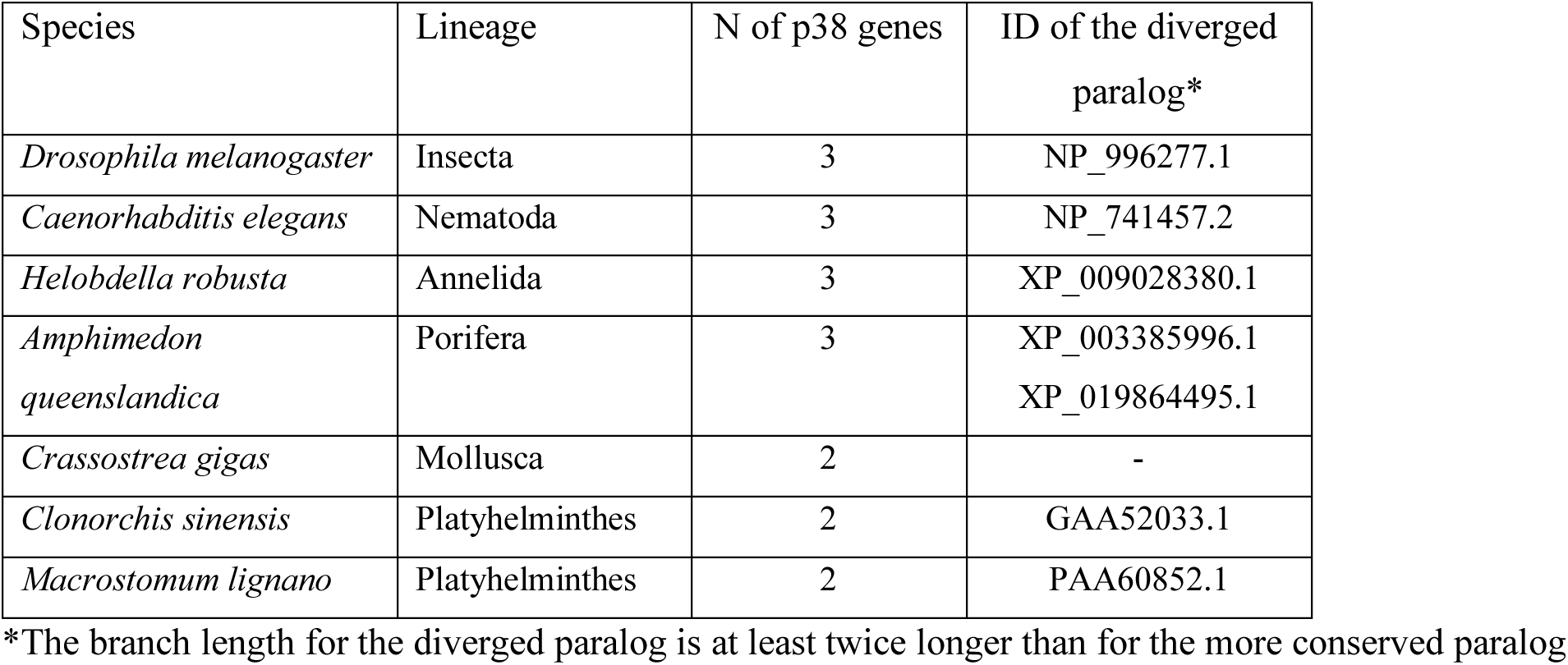
Number of p38-like paralogs in invertebrate species.

| Species | Lineage | N of p38 genes | ID of the diverged paralog* |
| --- | --- | --- | --- |
| *Drosophila melanogaster* | Insecta | 3 | NP_996277.1 |
| *Caenorhabditis elegans* | Nematoda | 3 | NP_741457.2 |
| *Helobdella robusta* | Annelida | 3 | XP_009028380.1 |
| *Amphimedon queenslandica* | Porifera | 3 | XP_003385996.1<br>XP_019864495.1 |
| *Crassostrea gigas* | Mollusca | 2 | - |
| *Clonorchis sinensis* | Platyhelminthes | 2 | GAA52033.1 |
| *Macrostomum lignano* | Platyhelminthes | 2 | PAA60852.1 |

\*The branch length for the diverged paralog is at least twice longer than for the more conserved paralog

Interestingly, p38γ-p38δ and JNK show longer branches, indicating a higher level of divergence, which is in agreement with Brewster et al. [27], who suggested that p38 and Hog1 retained more similarity with the ancestral protein than JNK. Furthermore, in this study, we have identified a fifth p38 paralog in vertebrates that clusters together with p38γ and p38δ. From now on we will call it p38γ/δ.

### p38 evolution in vertebrates

To better understand the p38 paralogs fate, we inferred a phylogenetic tree with an extended dataset for vertebrates. We included species relevant for our study due either to their big sizes (like whales and elephants) or their specific immune system (like bats or naked mole). Altogether, we collected 294 protein sequences from 63 species (Table S3, Supplementary). The tree (Fig. 2) showed that the evolution of p38 kinases in animals followed the scenario of the two rounds of whole genome duplication (WGD) in vertebrates and an additional lineage-specific WGD in bony fishes and Xenopus (Table 2). An exception is the group Cyclostomata which in our study is represented by two species *Petromyzon marinus* (has two paralogs, one is a sister branch to p38α-p38β, another is a sister branch to p38γ-p38δ) and *Lethenteron camtschaticum* (has only p38α and p38β). It is now thought that this group underwent two rounds of whole genome duplication as all other vertebrates. However this was an open question for some time because genomes of these animals exhibit the phenomena of asymmetric gene repertoire [28]. Our observation is likely the confirmation of this phenomenon, i.e. the specific pattern of genes retention and loss. However, the fact that *P. marinus*’s proteins cannot be assigned to any of the p38 vertebrate paralogs is unusual and illustrates how entangled can be paralogs evolution in some lineages. We also observed a distinct trend in teleost fishes, where only additional copies of p38α and p38γ were retained, but not of p38β and p38δ. The genomic location of the corresponding genes (they are distributed over two chromosomes in pairs p38α-p38δ and p38β-p38γ [2]), suggest that this is a selective retention rather than a random loss of the chromosomal segment. *Salmo salar* has four p38γ genes probably due to the additional round of WGD in salmonids.

**Figure 2.**
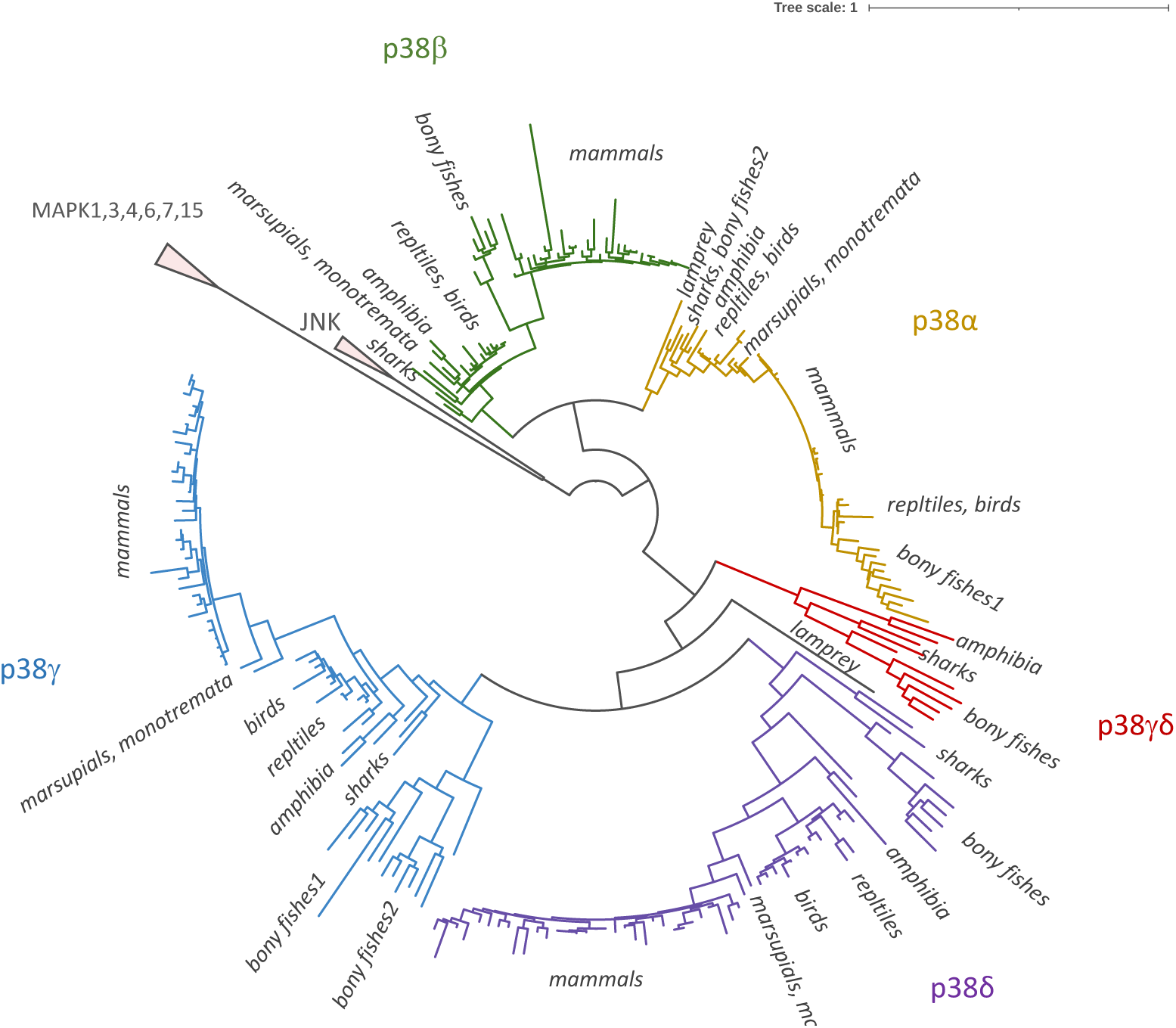
Maximum likelihood phylogenetic tree of p38 with an extended dataset for vertebrates. A new p38γ/δ paralog is highlighted in red.

**Table 2.**
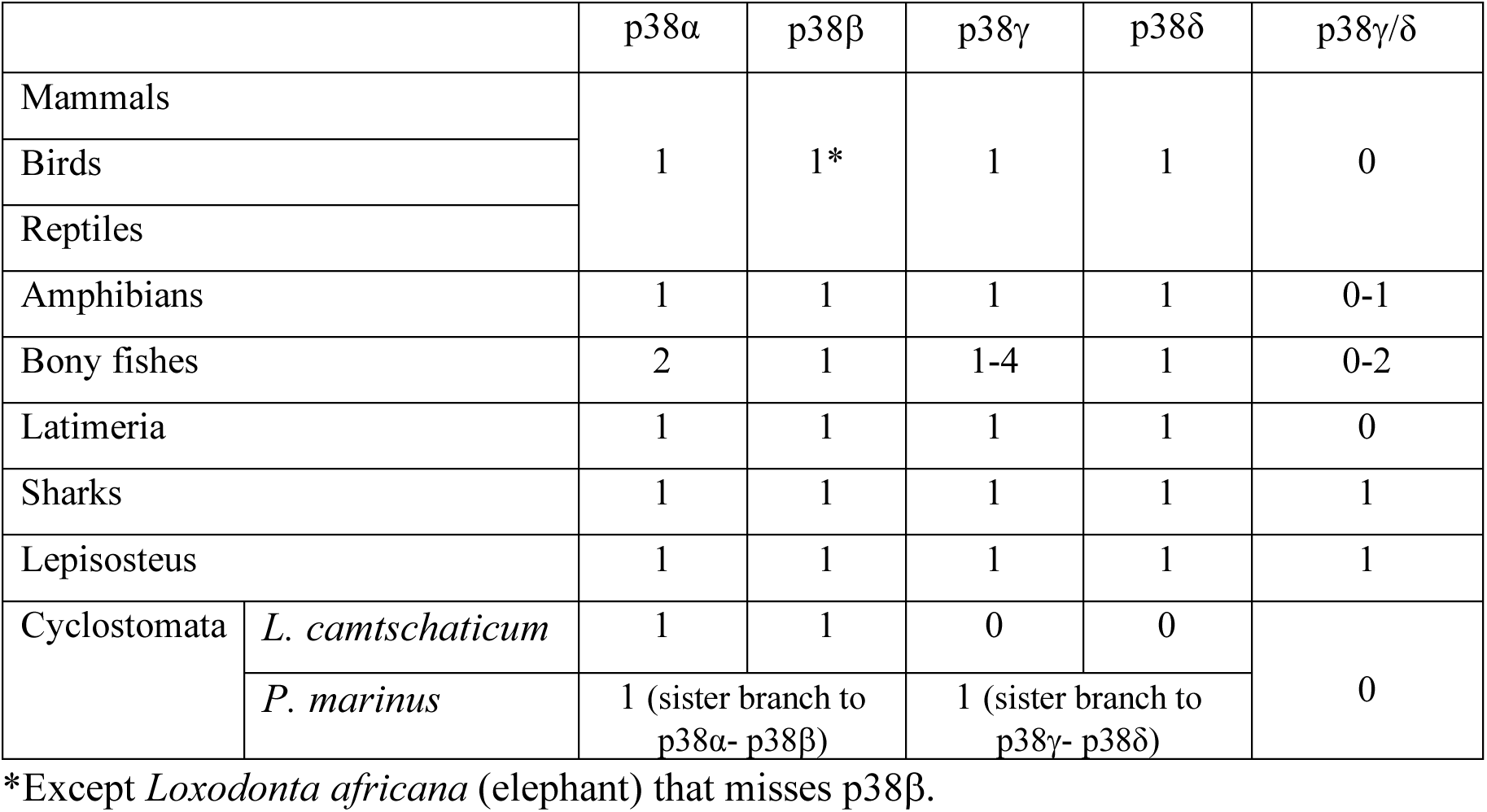
p38 paralog distribution in animal groups.

Another interesting finding is the new fifth paralog p38γ/δ present in amphibians, bony fishes, sharks, and a representative of ray-finned fishes *Lepisosteus oculatus*, but absent in mammals, birds, and reptiles (Tab. 2). p38γ/δ has most of the kinase active sites conserved (Fig. S1) and likely participates in p38-mediated stress response as other p38 paralogs. However, its absence from mammals, the most popular and studied group of animals, made it unnoticed until now.

It is possible that p38γ/δ may be a lineage-specific duplication. However, considering its diverse distribution in distinct animal taxa, it is most likely that p38γ/δ was lost from mammals, birds, and reptiles. By comparing the synteny of all five p38 paralogs, we concluded that p38γ/δ likely originated from a duplication of the p38β gene. Indeed, the position of plexin-b2 gene upstream of p38β is conserved in animals [6] and similarly plexin-b1 gene is located upstream of the p38γ/δ, although in the reverse direction (Fig. 3). Plexin b1 and b2 genes are close paralogs and their evolution is intertwined, alike that in the p38 family.

**Figure 3.**
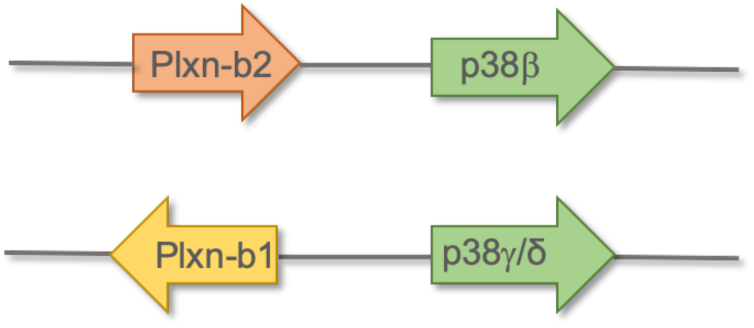
Schematic representation of synteny conservation for p38β and p38γ/δ and plexin genes in animals

As discussed in the previous section, p38γ and p38δ tend to have longer branches (Fig. 1), which suggests that these p38 paralogs underwent evolution with a higher mutational rate. To further test this hypothesis, we performed an analysis for positive selection using the codeml program from the PAML package. We used a tree with only mammalian sequences to avoid any possible artefacts due to the dataset bias. The results indicated the possibility for positive selection for p38γ (*p*-value based on *Χ*^2^ distribution is 0,02) and p38δ (*p*-value based on *Χ*^2^ distribution is 2,65×10^−6^), but neither for p38β nor for p38α. Strikingly, amino acids that contributed to positive selection play the role in the p38δ substrate binding site and in the hinge site of domain rotation during kinase activation (numbers 5 and 7, Fig. 4). This might indicate that p38γ and p38δ were retained in the mammalian genomes not only as the “back-up” for p38α, but also acquired individual features that could allow binding to specific partners.

**Figure 4.**
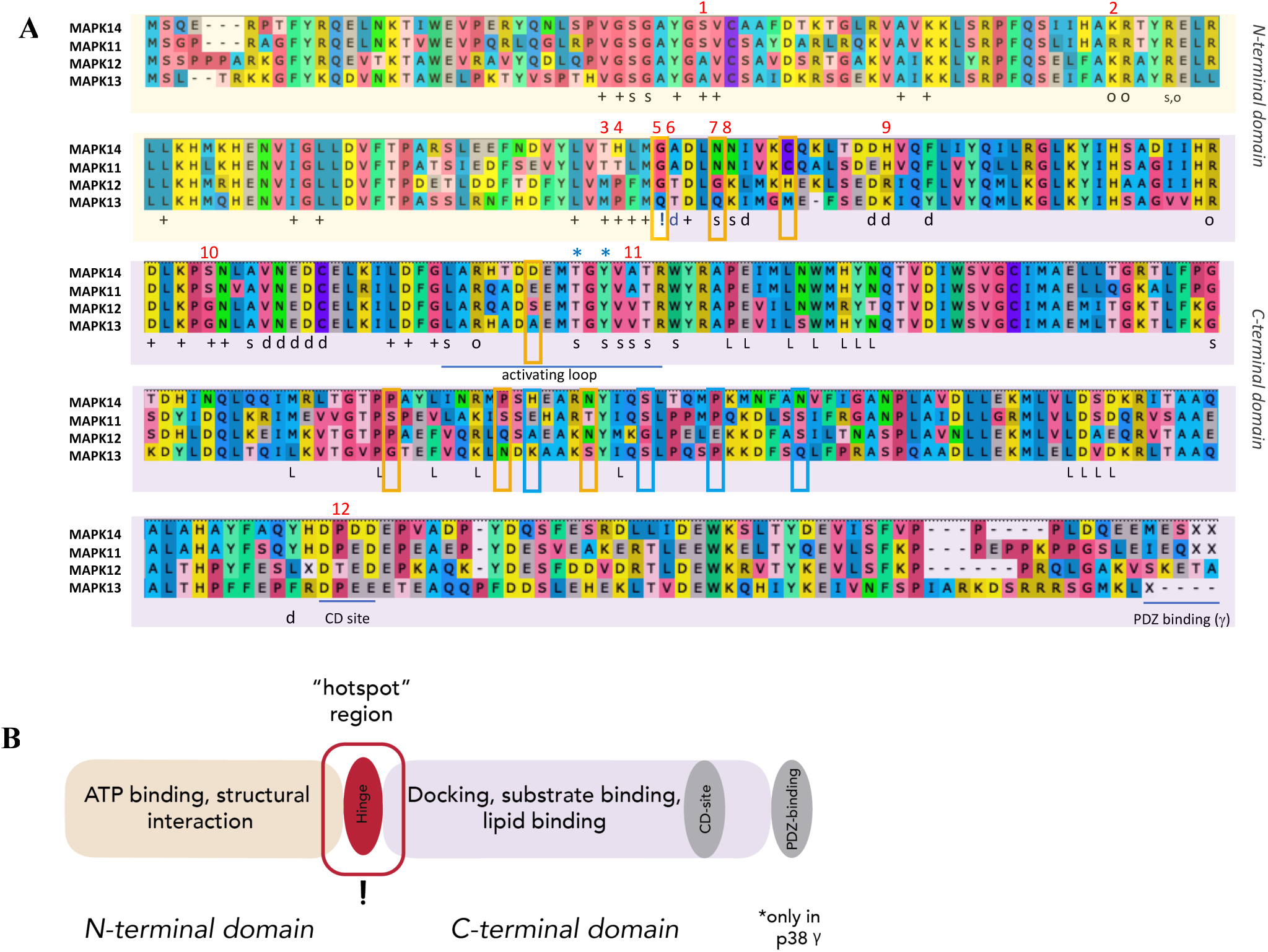
A. Alignment of the p38 consensus sequences in mammals. Designation: (+) ATP binding site; (s) substrate binding site; (o) AA interacting with the conserved threonine when phosphorylated, this is necessary for the folding into an active state conformation; (d) docking site, (L) lipid binding site; (!) hinge point, the two domains rotate around hinge point during activation; asterisks mark the phosphorylation sites specific to the stress MAPKs. PDZ binding domain is conserved only in p38γ. Red numbers point at the AA substitutions in functionally important sites. Blue frames highlight positively selected sites in p38γ, orange in p38δ. B. Schematic of the p38 protein domain architecture. The evolutionary hotspot region around the hinge point is highlighted.

In general, all four p38 kinases demonstrated high sequence conservation, even in the species where we expected to see differences (naked mole, bats). Nonetheless, we noticed a trend of N-terminal truncation in p38β and p38δ in the common minke whale *Balaenoptera acutorostrata* and in p38γ of the African bush elephant *Loxodonta Africana*. In addition, the genome of *L. africana* lacks the gene coding for p38β. This might be a result of the genome annotation inaccuracies, however, the p38α sequence in both species is complete and conserved, including its N-terminal end. This latter fact might support both the assigning p38α as the “main” paralog and suggest that the high weight animals have, indeed, undergone p38 paralog-specific sequence diversification and paralog loss. However, this hypothesis should be tested in further studies.

### Paralog diversification and evolution in mammalian p38s

To understand whether different p38 paralogs could exhibit distinct activities in the cell, we studied paralog sequence individuality in mammals. For this, we searched for paralog specific amino acid (AA) substitutions that could indicate specific functions. Our data show that most of the functional sites are highly conserved in all four paralogs, except the ATP-binding and the docking regions around the hinge point (after activation by dual phosphorylation the N-and C-terminal domains of p38 rotate around the hinge point). Interestingly, this region seems to be a “hotspot” for paralog diversification and likely affects each paralog’s substrate specificity and ATP binding efficiency. We highlighted 12 amino acid substitutions as the most probable to affect protein conformation and/or function (Fig. 4). Half of these sites are conserved between p38 subgroups, i.e. p38α and p38β share one variant and p38γ and p38δ share a second variant. The other half of the sites differ between all paralogs.

The most intriguing sites lie within the hinge point of p38δ (number 5: G->Q) and a nearby site involved in substrate binding (number 7: N->Q), both of which were detected as sites of positive selection by the codeml analysis. Although these amino acid changes are semi-conservative in its chemical nature and should not disrupt protein folding, the substitution of the conserved glycine can result in changes in electrostatic interactions, for example, interactions with phosphate groups [29]. In addition, due to its conformational flexibility, glycine is often present in structural turns where its substitution to any other amino acid is disfavored. Similarly, N->Q change can alter kinase activity in specific structural 3D contexts [29]. p38γ also carries a substitution in substrate binding site (number 7: N->G) which generally is considered neutral but can influence the activity in some kinases. Thus, we can suggest that these AA substitutions within the hotspot region may be the basis of p38 paralog specific roles in the cell. Possible effects of the 12 highlighted amino acid substitutions are summarized in Table 3.

**Table 3.** Paralog specific amino acid substitutions in p38 functional sites.

| Number on Figure 3 | Substitution | P38 paralog * | Associated function | Possible effect | Reference |
| --- | --- | --- | --- | --- | --- |
| 1 | S → A | p38γ and p38δ | ATP binding | Can alter orientation of ATP in the binding site or optimal pH range for the enzyme activity | [31] |
| 2 | K → R | p38β | Conformational changes during activation | Mostly neutral, can be non-conservative in specific tertiary structures | [32]–[34] |
| 3 | T → M | p38γ and p38δ | ATP binding | Can alter kinase activity as was shown for RET tyrosine kinase | [35] |
| 4 | H → T<br>H → P | p38β<br>p38γ and p38δ | ATP binding | Histidine is efficient proton exchanger in protein active sites; mutation can change binding specificity. | [29], [36] |
| 5 | G → Q | p38δ | Hinge site | Can influence interactions with other AA or phosphates; substitution of G in turns is rare | [29] |
| 6 | A → T | p38γ and p38δ | Docking site (specificity to substrate) | Can alter AA orientation in the docking site | [29], [31] |
| 7 | N → G<br>N → Q | p38γ<br>p38δ | Substrate binding | Can change kinase activity or binding efficiency | [29] |
| 8 | N → K | p38γ and p38δ | Substrate binding | Can alter binding efficiency | [37] |
| 9 | H → R<br>H → K | p38γ<br>p38δ | Docking site (specificity to substrate) | Can alter binding efficiency | [29], [38], [39] |
| 10 | S → G | p38γ and p38δ | ATP binding | Can alter binding specificity | [40] |
| 11 | A → V | p38γ and p38δ | Substrate binding | Both AAs are little reactive, but can act in substrate recognition differently. V is less conformationally flexible than A, especially in α-helices | [29], [41] |
| 12 | P → T | p38γ | CD site | Substitutions of P are rarely tolerated when in specific protein folds. P->T mutation was shown to influence activity of several enzymes if happens in active sites. | [42]–[44] |

\*relative to reference paralog p38α

Overall, the variation in the hotspot region can be defined as mild, i.e. substitutions happen between AA with similar biochemical characteristics. Therefore, such substitutions are not expected to inhibit or alter the kinase’s function. At the same time, they have a potential to modulate efficiency of ATP binding, substrate binding specificity and affinity in each p38 paralogous protein. In this way, each paralog would be tuned to its specific target or regulator in particular cellular pathways or contexts. Future experiments using site-directed mutagenesis could show if the differences in the near-hinge site region can be indeed used to characterize each p38 paralog’s individuality. At least, one such experiment has been already done, where changing the threonine within the ATP-binding site (number 3 on Fig. 4A) of p38α and p38β altered the sensitivity of these kinases to an ATP-binding inhibitor [30].

Next, we searched for conserved substitutions in the fifth p38γ/δ paralog, which is absent in mammals. Sequence comparison of p38γ/δ to mammalian p38 paralogs revealed that amino acids within the ATP binding site are conserved between p38γ/δ and p38γ and p38δ (Figure S1, Supplementary). At the same time, it has unique substitutions within the hotspot site: in the docking and substrate binding regions (numbers 4-9 on Figure S1). This shows that all five paralogs can be distinguished based on the comparison within this region. For example, the A->Q substitution in the site associated with the docking function (number 5 in Fig. S1, homologous to number 6 in Fig. 4A) where alanine, generally non-reactive and hydrophobic, is changed to polar glutamine with a large side chain. Similarly, histidine (positively charged) is changed to isoleucine (uncharged, hydrophobic), which is a non-conservative change and can likely affect docking specificity or strength (number 9 on Fig. S1 and on Fig. 4A). p38γ/δ also has conserved substitutions of the two docking AAs outside of the hotspot, unlike other p38 paralogs (numbers 12 and 15, Fig. S1). This suggests that p38γ/δ may have evolved the capacity to interact with new substrates and/or participate in different pathways and respond to new stimuli. Further experiments in understanding the cellular functions of p38γ/δ and its role in stress response will provide interesting insights into the role of the p38 family in animal physiology.

Our consensus comparison approach can be used for analyzing sequence evolution in any other p38 homologs. For example, we looked at the invertebrate p38s. Surprisingly, they, too, exhibit high sequence conservation, especially in the docking site, in the amino acids surrounding TGY motif, and among amino acids involved in the structural interaction with the phosphorylated threonine of the TGY motif. However, invertebrate p38 show higher diversification within the ATP-binding site, especially the p38 proteins that cluster with p38γ and p38δ.

Sequence comparison analysis of animal p38 kinases shows that all paralogs have lineage-specific AA substitutions. Many of these substitutions are positioned at the surface of the folded p38s as predicted by 3D structure analysis in I-TASSER (Table S4, Supplementary). Therefore, these substitutions can be potential candidates for further functional studies of the p38 kinases. A particular case is the PDZ binding region that is conserved in p38γ kinase in all the vertebrate analyzed here but not present in any other paralogs. This domain is involved in protein interactions and might be important for p38γ functions by forming different protein complexes. Interestingly, lipid binding sites that have been described based on the crystal structure of p38α kinase [22] are well preserved in all five p38 kinases in all lineages with a rare exception (in vertebrates, out of 16 positions 9 are identical and 5 are changed to the same chemical AA type). This suggests the importance of the lipid binding in p38 regulation and, therefore, highlights the need of future functional studies in this direction.

### Paralog specific amino acid motifs of p38/Hog1-like proteins in unicellular relatives of animals

The uncertain position of the p38-like proteins of unicellular relatives of animals in the phylogenetic tree does not allow us to discern whether those homologs are “animal-like” p38 kinases or “fungi-like” Hog1. Since the binding activity of proteins is often defined by just one or a few amino acids, changes in active sites are difficult to trace in a phylogenetic tree. Therefore, we performed additional sequence comparison analysis, now including sequences from choanoflagellates, filastereans, corallochytreans, and ichthyosporeans in order to determine whether they can be assigned to one of the p38, JNK or Hog1 groups (Fig. 5). Strikingly, all homologs from the unicellular organisms preserve the TGY phosphorylation motif characteristic to p38 and Hog1. This confirms that these proteins belong to p38/Hog1 kinases and distinguishes them from other MAPKs. Interestingly, p38-like proteins from unicellular relatives of animals share high similarity with both animal and fungal stress kinases in several key functional sites (Fig. 5). Lipid binding sites show low level of conservation as compared to animals, with only half of the sites conserved.

**Figure 5.**
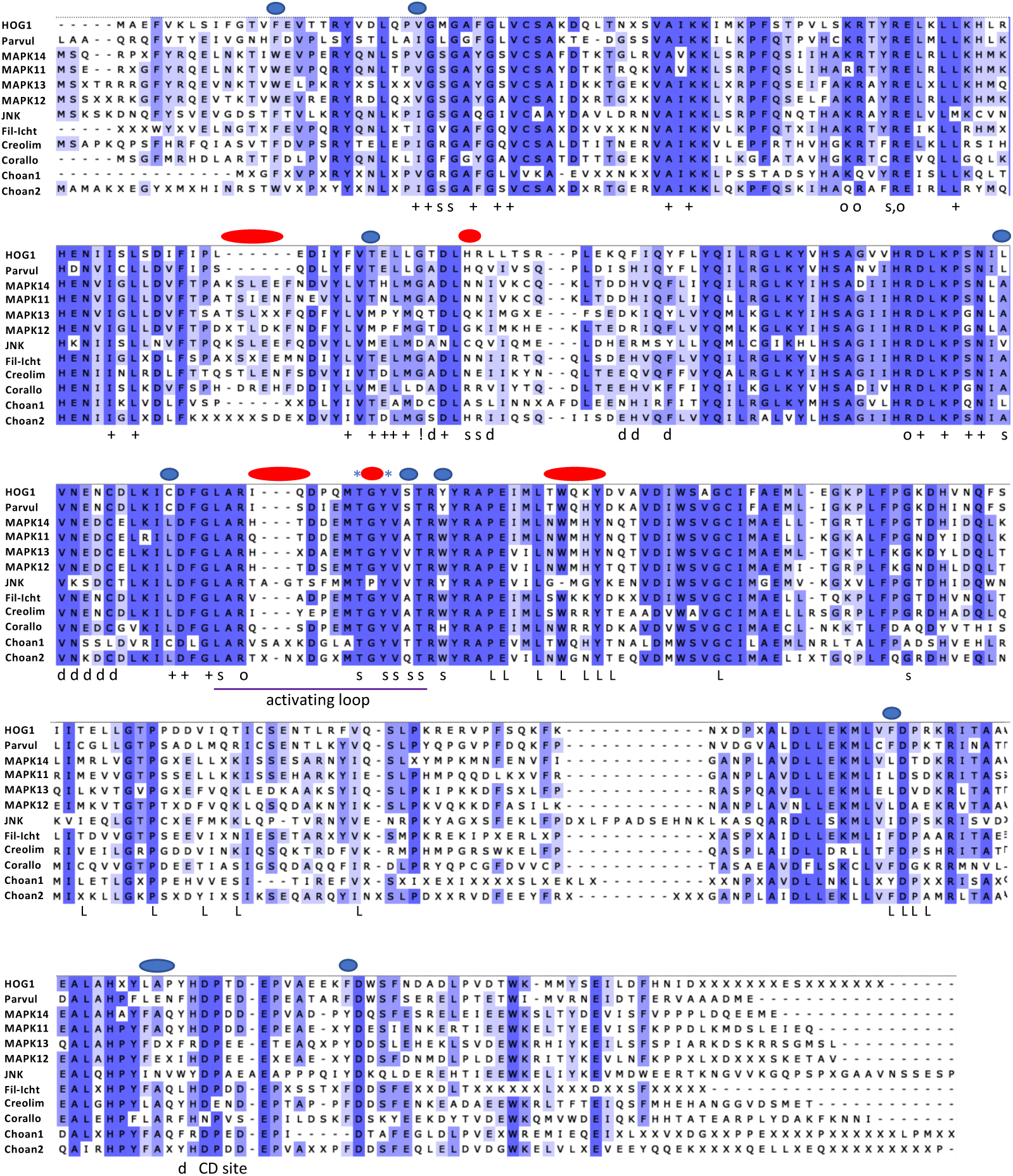
Alignment of the consensus sequences of p38, JNK, HOG1 and the homologs from unicellular opisthokonts. Alignment color-code correspond to AA identity, from the highest (dark blue) to low (white, less than 40% of positions in a column is conserved). Designation: (+) ATP binding site; (s) substrate binding site; (o) AA interacting with the conserved threonine when phosphorylated, necessary for folding into active state conformation; (d) docking site, (L) lipid binding site; (!) hinge point, the two domains rotate around hinge point during activation; asterisks mark the phosphorylation sites specific to the stress MAPKs. Blue circles mark sites shared by two or more paralogs, red circles mark regions that can distinguish between p38, JNK, and Hog1. Parvul – sequence from *P. atlantis*, Fil-Icht – consensus sequence for the clade filasterans+ichtchyosporeans from the tree on Figure 1, Creolim – sequence from *Creolimax fragrantissima*, Corallo – sequence from *Corallochytrium limacisporum*, Choan1 and Choan2 – consensus sequences from choanoflagellates choano-1 and choano-2, respectively.

Overall, the sequence similarity analysis shows mosaic conservation between p38, JNK, Hog1, choanoflagellate choano-1 and choano-2, and filastereans-ichthyosporeans-Corallochytrium (f-i-C). I.e., homologs from the unicellular relatives of animals share similarities with p38 in some sites and with JNK or Hog1 in others (blue ovals in Fig. 5). Such pattern implies that all stress activated kinases evolved from an ancestral protein that originated before the split between Holozoa (metazoans and their closest unicellular relatives) and Holomycota (fungi and their closest unicellular relatives). Moreover, functional diversification of all p38 kinases possibly occurred due to amino acid substitutions in the substrate binding and docking sites in the near-hinge site (the hotspot), since the variation within this region is increased in the p38 kinases from the unicellular holozoans, as well as in animals.

It is likely that the f-i-C group, the animal p38, and the fungal Hog1 group diverged less from the ancestral kinase, than the JNK and both choanoflagellate clusters. The evolutionary distance between the choanoflagellate choano-1 and animal p38 kinases is the longest among all comparisons, except with Hog1 (Table 4). In agreement with the tree on the Figure 1, sequence analysis also indicates that choanoflagellate choano-2 shares more similarities with p38 rather than with Hog1. Because JNK stands out from the whole group as the most diverged paralog, we propose to call the orthologs from the unicellular holozoans p38/Hog1-like kinases.

**Table 4.**
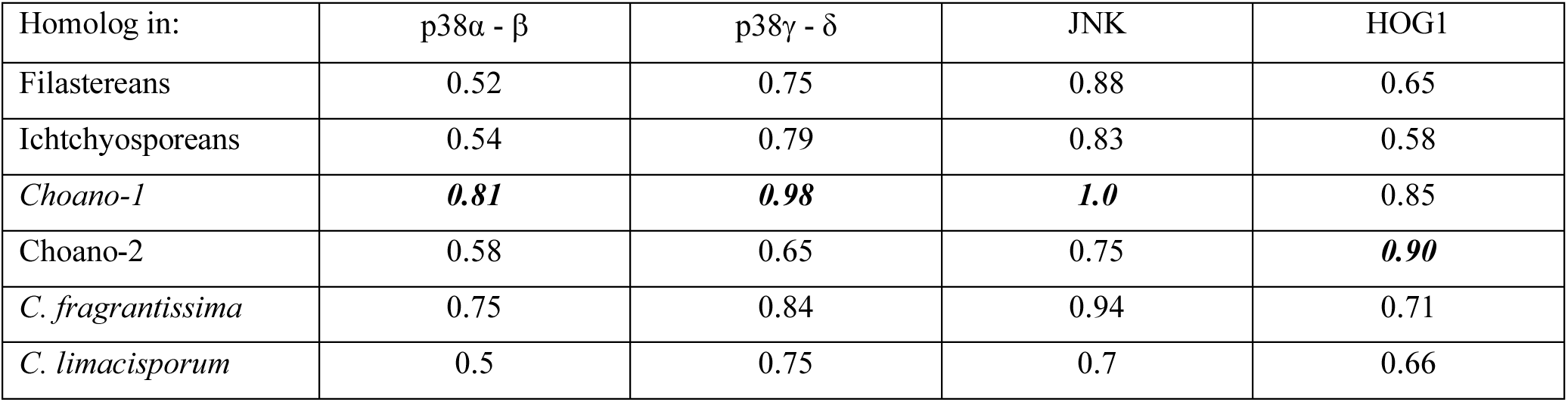
Evolutionary distances between different clusters of p38 and p38-like proteins. Choano-1 from choanoflagellates is the most divergent from the unicellular holozoans. Choano-2 has the longest distance to Hog1.

### p38/Hog1 protein of *Capsaspora owczarzaki* reveals functional similarity to the fungal Hog1

To better understand the origin and evolution of the stress-activated kinases, we performed stress studies on *C. owczarzaki* cultures. The filasterean *C. owczarzaki* is one of the most studied unicellular relatives of animals. Its p38-like homolog clusters within the f-i-C group on the tree (Fig. 1) and shares sequence similarities with p38, as well with Hog1. It is known that all three types of kinases – p38, JNK, and Hog1 – can be activated by osmotic shock and it has also been shown that expression of p38 and JNK can rescue Hog1 knockout in yeasts [5]. Therefore, we subjected aggregative *C. owczarzaki* cells to osmotic shock by exposing them to 0.1 M NaCl concentration in the culture medium. Next, we used the qPCR technique to measure mRNA expression levels of six *C. owczarzaki*’s genes, whose homologs in animals or fungi are known to regulate different types of stress response. In particular, we selected glycerol-3-phosphate dehydrogenase (GPD1), Heat shock protein 90 (Hsp90), MYC, apoptosis-inducing factor (AIF), NF-κB, and the p38/Hog1-like homolog identified in this study (further called COp38/Hog1).

In this stress condition, *C. owczarzaki*’s cells seem to activate a pro-survival mechanism by downregulating the expression of *AIF* mRNA after 45 minutes after the stress induction and upregulating the expression of *Myc* and *NF-κB* mRNA after 3,5 hours (Fig. 6A). We also observed an increase in the expression of *GAPDH* mRNA after 45 minutes post-induction. *GAPDH* gene is commonly used as a house-keeping gene, but its regulation was affected by stress, which is in agreement with the previously published studies [45]–[47].

**Figure 6.**
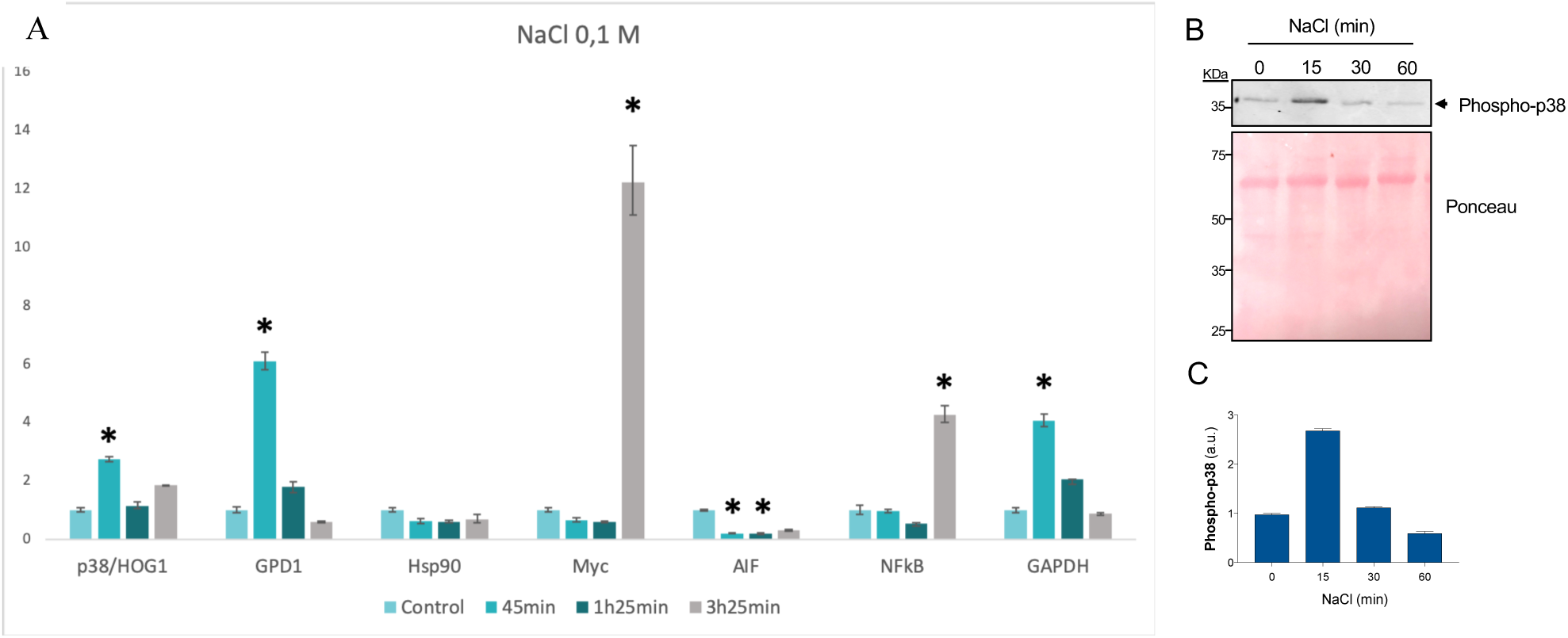
(A) mRNA levels (fold change) measured in *C. owczarzaki* cells by qPCR conditions after 45 minutes in osmotic stress, 1 hour 25 minutes, and 3 hours 45 minutes of incubation. Error bars indicate standard error (see Methods). Asterisks mark significant change in the mRNA expression (p-value < 0,05). (B) Osmotic stress induces the phosphorylation of COp38/Hog1. Adherent *C. owczarzaki* cells were incubated for the indicated times with 0.1 M NaCl. 5 µg of total cell extracts were immunoblotted with anti-phospho-p38 antibody. Top panel shows representative blots from two experiments. Bottom panel shows Ponceau staining of the membrane before immunoblotting, as a protein loading control. The position of molecular weight markers in KDaltons is indicated. (C) Phospho-p38 band intensities in (B) were quantified using the Odyssey infrared imaging system and represented as histograms. Values are given as mean ± SEM from two representative gels in duplicates. a.u.: arbitrary units.

Intriguingly, *COp38/Hog1* and *GDP1* mRNA expression was also upregulated after 45 minutes of incubation under the osmotic stress conditions and decreased at 85 min (Fig. 6A). Upregulation of GPD1 downstream of Hog1 is specific to fungi in the osmotic stress conditions [27]. GPD1 is involved in Sln1-regulated pathway, which in yeasts mediates osmotic stress response and results in synthesis of glycerol. By accumulating glycerol, fungal cells compensate for the increased salinity in the environment. Accordingly, *C. owczarzaki* seems to have a fully conserved Sln1-regulated pathway (Table S5, Supplementary). Together with our results, this suggests that the species might exhibit an osmotic stress response using a mechanism similar to that described in fungi. Although upregulation of p38 mRNA expression has not been shown in animal stressed cells, we cannot rule out that this mechanism is present in *C. owczarzaki* either as species specific or ancestral feature.

Cellular stress response in eukaryotes is mediated by signaling cascades that are activated by the sequential phosphorylation of several kinases. In order to investigate whether phosphorylation of COp38/Hog1 is also involved in the regulation of the osmotic stress, we stimulated *C. owczarzaki* adherent cells with 0.1 M NaCl. At different times post-induction, cells were collected and protein extracts obtained. The phosphorylation of COp38/Hog1 was evaluated by western blot, using phospho-specific antibodies. These antibodies recognize the conserved TGY motif when phosphorylated, i.e. they recognize p38 kinases in their activated state. As shown in Figure 6B and 6C, a phosphorylated 37kD protein, corresponding to COp38/Hog1, was detected. Phosphorylation of p38/Hog1 was significantly increased at 15 min post-induction compared to control cells, but decreased after 30 min. This pattern of quick phosphorylation and subsequent dephosphorylation of p38 or Hog1 is crucial and typical of the stress response in other organisms and suggest that COp38/Hog1 is, indeed, mediating such a function in *C. owczarzaki* [48], [49]

None of the available antibodies against total p38α, p38γ, p38δ, or phospho-JNK detected any protein of the expected size in *C. owczarzaki* cell extracts. These antibodies were raised against mammalian proteins and it is possible that they failed to recognize the distant p38 homolog from *C. owczarzaki*. In contrast, the phospho-specific p38 antibody, recognizes the small phosphorylated TGY motif, which are very well conserved across species. It is important to note that for the analysis of COp38/Hog1-like kinase phosphorylation under osmotic stress, we first used aggregative *C. owczarzaki* cells. However, aggregative cells showed high level of COp38/Hog1 phosphorylation even in unstressed conditions (control cells not exposed to increased NaCl concentration), which is in agreement with the results of the phospho-proteomics analysis of *C. owczarzaki*’s life stages described in [50].

The results described here show that the phosphorylated p38/Hog1 kinase in *C. owczarzaki* can be detected with antibodies raised against the mammalian phosphorylated protein and that its activation by osmotic stress follows a pattern of phosphorylation-dephosphorylation similar to other eukaryotic cells [13]. In combination with qPCR expression studies, we demonstrate that *C. owczarzaki* is likely to exploit fungi-like stress response to osmotic shock. In addition, structural comparison of the modelled 3D structure of COp38/Hog1 to Hog1 and p38α as well indicates its higher similarity to yeast Hog1 (RMSD value is 1,9 and Z-score is 45) rather than to p38α (RMSD value is 2,4 and Z-score is 41,1). Similarly, it was suggested previously that some physiological and morphological features of the unicellular relatives of animals could be acquired from the common ancestor of animals and fungi [51].

## Discussion

In this study, we pinpoint the origin of p38 stress kinases and their consequent evolution in animals. We also describe the molecular basis behind p38 paralogs individuality. In particular, we demonstrate that stress-activated kinases originated in the common ancestor of Opisthokonts (the group comprising Metazoa, Fungi, and their unicellular relatives). The ancestral p38/Hog1 protein evolved into p38 and JNK in animals and into Hog1 in fungi. The homologs from extant unicellular relatives of animals (choanoflagellates, filastereans, and teretosporeans) show similarity to both p38 and Hog1 proteins, likely mirroring the features of that ancestral p38/Hog1 protein. Interestingly, the genomes of choanoflagellates contain one Hog1-like paralog and one p38-like paralog, which might be a unique feature of these organisms. Strikingly, the proteins from the unicellular relatives of animals share high similarity with animal p38s and Hog1 within functionally important regions (TGY phosphorylation motif, ATP binding site, activation loop with the associated substrate binding sites). They also preserve five amino acids (lysine and four arginines) that have been shown to pull the kinase’s structure into the active conformation upon the dual phosphorylation event [52]. Thus, we can assume that these newly identified p38/Hog1-like orthologs perform functions similar to the animal and/or fungal stress kinases. Indeed, we showed that the dynamics of activation of p38/Hog1-like kinase in *C. owczarzaki*, a unicellular relative of animals, follow the pattern observed in animals and fungi under hyperosmotic stress. Increased mRNA of GPD1 is, however, resembling the signaling pathway in yeasts. *C. owczarzaki* is, therefore, an interesting example of an organism from an intermediate taxonomic position between Metazoa and Fungi likely preserving physiological features both from the fungal ancestor and the unicellular ancestor of animals. Answering whether p38/Hog1-like kinase in *C. owczarzaki* and other unicellular relatives of animals indeed participate in stress response and whether they are involved in pathways more related to fungi or animals, will unravel how the stress response network evolved in animals.

Therefore, based on this and supported by previous studies (see review [27]), we hypothesize that the common ancestor of Holozoa (animals and their unicellular relatives) possessed at least one MAPK-like stress kinase with functions involved in response to osmotic stress. However, it could participate in the general stress resistance of the cell. Detection of and response to environmental changes in salinity is an ancient function serving the cell’s integrity. Therefore, it is not surprising that the elements involved in its regulation are conserved between fungi and holozoans. For example, it has been shown that p38 from animals can regulate osmotic physiological response in yeasts [27]. During evolution in the animal lineage, rounds of WGD would allow expansion of gene families involved in the stress response and diversification of cells’ reaction to a multitude of stress stimuli in different cell types. As a result, vertebrates have 4-5 p38 kinases and 3 JNK. While these are very similar paralogs, they still demonstrate specificity towards inhibitors and substrates at least to some extent [13]. Our analysis identified the probable evolutionary hotspot where mutations can lead to paralog specific activity. While substrate binding sites in the activating loop are conserved in all animal paralogs and also in p38/Hog1-like orthologs from unicellular holozoans, the amino acids involved in substrate docking and substrate binding around the hinge site show variation between the paralogs. We showed that amino acid substitutions within the hotspot are semi-conservative and could only be responsible for mild changes in the local charge or structure. This would allow keeping the overall function of the enzyme while modulating substrate specificity or binding efficiency, since protein binding sites are usually defined by only a few amino acids and small allosteric changes [53]. Indeed, substrate docking in p38s was shown to be responsible for their specificity and efficiency in different cellular cascades [20]. Accordingly, it was experimentally shown that mutation of just two amino acids within the docking site could change substrate specificity of p38 to that of ERK and vice versa [54]. Therefore, the sequence evolution of p38 paralogs probably enabled the cell to enrich in a variety of responses to different stresses and in different cell types in animals. This would also enhance precision in the regulation of the kinase activity and increase the number of interacting proteins in the pathway, as similarly discussed in [2], [6], [27], [55], [56]. Such functional diversity could also have led to a very complex crosstalk between p38 and JNK kinases in different conditions and different cell types, as was shown in some studies [57].

Paralog diversification in the p38-JNK group in animals can be seen as a classical example of the phenomena where one copy (p38α) preserves the original function and other copies have more flexibility and undergo further divergence (p38γ, p38δ, JNK, and the newly described here p38γ/δ). The evolution of p38γ, in particular, resulted in the acquisition of a new binding site for PDZ-containing proteins, such as α1-syntrophin, SAP90/PSD95 and SAP97/hDlg. This binding site is essential for the phosphorylation of these proteins by p38γ under stress conditions [21], [58], [59]. We also described a new, fifth p38 paralog, p38γ/δ, present in several animal lineages. This protein is characterized by specific amino acid substitutions within the hotspot region, indicating that it might have a specific set of targets. This finding further demonstrates the complexity of p38 signaling in animals.

A similar strategy of paralog duplication and diversification can be seen in some invertebrates. For example, *C. elegans, D. melanogaster* and a few other species underwent lineage-specific duplication, and one of their paralogs is characterized by higher sequence divergence. *Drospophila*’s p38c, for example, even lost the TGY motif typical to p38. Experiments with other long branched paralogs described in this study will bring more understanding of p38 evolutionary history and the capacity of paralogs to acquire new characteristics. Our detailed evolutionary scenario presented here is a valuable resource for further research on the p38 family. In particular, it can help in identifying amino acids that might be relevant to understanding p38 paralog individual roles in diseases and to discovering new functionally important sites that novel therapeutic inhibitors can target.

## Methods

### Data collection and phylogenetic analysis

Protein sequences were collected by running blastp and tblastn search against protein database at NCBI and GenBank nucleotide database, respectively, with human p38 proteins as query (NP_002745.1, NP_001306.1, NP_002960.2, NP_002742.3). Sequences from Breviatea were obtained from [60], CRuMs from [61], and *S. destruens* from [51]. Hits with more than 30% identity and 75% query coverage were selected. Incomplete and misassembled sequences were manually corrected. A multiple sequence alignment was built in MAFFT [62], and the alignment was manually checked and trimmed using Ugene program [63]. The phylogenetic tree of Figure 1 was inferred using IQtree [64] with the model LG+G4+F and 1000 ultrafast bootstrap replicates to test nodal support. The tree of Figure 2 and was inferred using RAxML [65] with the PROTGAMMALG model as chosen by performing a RAxML parameter test. Tree visualization was done in iTOLtree [66]. All trees in Newick format and all multiple sequence alignments can be found in the Supplementary Files 1-3 (can be accessed at Figshare DOI: <u>10.6084/m9.figshare.19487627</u>).

### Test for positive selection

To test whether any of the p38 paralogs in mammalian lineage underwent positive selection, we first inferred an ML tree with mammalian sequences using RAxML as described above. Next, we created an amino acid -nucleotide alignment using PAL2NAL software [67]. The test for positive selection was done using the codeml tool in PAML software [68]. We used site-branch model analysis to test each of the four branches with the following parameters: model = 2, NSsites = 2, fix_omega = 0, and omega = 1. The null hypothesis in this case was estimated under parameters: model = 2, NSsites = 2, fix_omega = 1, and omega = 1. For statistical assessment, twice the difference between the log likelihood of the alternative and the null (the ratio is the same for all branches) hypotheses were compared with the *χ*^2^ distribution. The *χ*^2^ distribution test for statistical support was used. Each estimation of alternative hypotheses was run twice by codeml with varying parameter “omega” to test for contingency. Parameter “cleandata” was set to 0, allowing for retaining gaps in the alignment. The input phylogenetic tree was built using RAxML as described earlier.

### Comparative sequence analysis

To identify paralog specific substitutions, we first built a consensus sequence using EMBOSS Cons using 7 protein sequences for each lineage. Multiple sequence alignment was done and manually corrected in Ugene. AA substitutions were highlighted based on the analysis of AA biochemical qualities and literature search. Functionally important sites were mapped according to the conserved domains map at Conserved Domain database at NCBI [69] and the following studies [22], [70]–[76].

### Synteny analysis

Gene surroundings of p38γ/δ were assessed by analyzing gene maps in the genomic viewer at NCBI available at Gene database within the region of 15 Kbp in the following species: *Xenopus laevis, Danio rerio, Lepisosteus oculatus, Rhincodon typus*.

### Calculation of the evolutionary distances

Between group mean distances were calculated in MEGAX [77], version 10.1.7, based on the tree T1 and the corresponding multiple sequence alignment using default parameters.

### Cell culture, RNA extraction, and qPCR experiment

*C. owczarzaki* cells (strain ATCC^®^30864) were grown axenically in the ATCC medium 1034 (modified PYNFH medium) with the addition of 10% (v/v) fetal bovine serum (Sigma-Aldrich, #F9665) at 23ºC. Cells were seeded in 12-well plates at the density of 2.4***Χ***10^6^cells/ml and placed at agitation at 50 rpm for aggregate formation as described in [78]. At the same time point, the osmotic shock was induced by adding 0.1 M of NaCl (Sigma-Aldrich #S1619). NaCl solution was prepared in the culturing medium. Whole RNA was extracted from the cells in triplicates for each condition after 0 min (control), 45 min, 1 h 25 min, 3 h 45 min of incubation using Trizol (Invitrogen/Thermo Fisher Scientific, #15596026) as described in [79]. Subsequently, the samples were treated with DNAseI (Roche, #4716728001) and purified using the RNeasy minikit, (Qiagen, #74104). The RT-PCR was done using SuperScript III First-strand synthesis system for RT-PCR (Invitrogen, #18080-051) with the random hexamer primers, following the manufacturer’s protocol. The resulting cDNA was treated with RNaseH (Invitrogen, #AM2293).

Primers for each gene for the qPCR reaction were designed in Primer3 Plus [80]. The primer sequences can be found in the Table S6, Supplementary. Primers’ specificity was tested by PCR amplification using the cDNA and by generating a standard curve with cDNA dilutions 1; 0.1; 0.01; 0.001; 0.0001; 0.00001 with the iTaq Universal SYBR Green Supermix kit (Bio-Rad, #172-5121). Using the same kit, qPCR was carried out with the iQ5 multicolor Real-Time PCR detection system with iCycler™ Thermal Cycler (Bio-RAD) using the Bio-RAD iQ5 software. All the conditions were tested in triplicates in addition to no-template control and a negative control with no-RT RNA. PGK1 was used as a housekeeping gene.

Data analysis and statistics test were carried with the program REST 2009 [81]. Fold change was calculated using the ΔΔCt method [82], [83]. PGK1 was used as a housekeeping gene.

### Defining NaCl working concentration

*C. owczarzaki* cells viability in hyperosmotic conditions was assessed by using the resazurin (Sigma, R7017) assay as following. We subjected the cells to the array of NaCl concentrations in the culturing media: 0.0125 M, 0.025 M, 0.05 M, 0.1 M, 0.2 M, 0.3 M, and 0.4 M. After 48h incubation we added resazurin reagent following the manufacturer’s protocol and measured the absorbance at 600 nm using fluorometer Infinite M200 (Tecan) after additional 48 h incubation with resazurin. Based on this data, 0.1 M NaCl concentration was chosen for hyper-osmotic stress experiments (Table S7, Supplementary). This concentration is in line with conditions for hyperosmotic stress experiments in mammalian cell cultures and fungi [84]-[86].

### Western Blot

Adherent cells were cultured as described earlier, and the osmotic shock was induced by adding 0.1 M of NaCl solution in the culturing media to the cells. At the indicated time points, the media was aspirated, and the lysis buffer was added directly to the cells. The cells were collected by pipetting up-and-down and the samples were immediately frozen in liquid nitrogen and stored at −70ºC until ready to proceed. Total protein extraction was performed as follows: samples were thawed on ice and sonicated on Digital Sonifier (Branson, model 102C) at the amplitude 15% with 15 sec pulse and 30 sec pause for 8 cycles on ice. The samples were then incubated on ice for 15 more minutes and centrifugated at 20000g, 20 min, 4ºC. The supernatants containing proteins were immediately frozen in liquid nitrogen and stored at −70ºC. Before freezing, aliquots were taken to measure protein concentration by the Bradford method (BioRad, #5000205) according to the manufacturer’s protocol. (Lysis buffer: 50 mM Tris pH 8, 150 mM NaCl, 0.1% SDS, 5 mM EDTA, 5 mM EGTA, 1% IGEPAL, 1mM MgCl_2_, 0.5 mM PMSF, 1 mM DTT, 1 mM Na_3_OV_4_, 1mg/ml pepstatin A (Sigma #P5318), 1 tablet of PhosStop (Roche, #4906837001) per 10 ml of the buffer, and 1 tablet of Complete-Mini (Roche, #4693116001) per 10 ml of the buffer).

Protein samples (5µg) were resolved by SDS-PAGE in 10% acrylamide gels and transferred to nitrocellulose membranes, which were blocked for 30 min with TBS-T Buffer (50 mM Tris/HCl, pH 7.5; 0.15 M NaCl; 0.05% (v/v) Tween) containing 10% (w/v) non-fat dry milk. The antibodies (0.5 µg/ml) were incubated overnight at 4°C, in TBS-T buffer with 5% (w/v) non-fat dry milk. Secondary antibodies, conjugated with a fluorochrome were diluted 1:5000 in TBS-T buffer with 5% (w/v) non-fat dry milk. Signal of the secondary antibody was detected with Odyssey Imaging system (Li-Cor). The antibodies used were anti Phospho-p38 (Thr180/Tyr182) antibodies (Cell Signaling #9211); anti Phospho-JNK (T183/Y185) antibodies (R&D Systems #MAB1205); anti p38α antibodies (Santa Cruz #sc-535); and secondary antibody Alexa Fluor 680 Goat Anti-Rabbit (Molecular Probes #A-21109). Anti-p38γ and -p38δ antibodies were raised and purified as described in [87].

### 3D structure reconstruction and structural comparison

The 3D model for COp38/Hog1 (Supplementary File 4, Figshare DOI: <u>10.6084/m9.figshare.19487627</u>) and surface residues was calculated in I-TASSER [88]. Structure comparison was done in DALI software [89]. For the comparison analysis the following models were used: human p38α crystal structure from the Protein Data Bank (https://www.rcsb.org/3d-view/5ETC) and the 3D model P32485 (HOG1_YEAST) from Swiss-Model for S. cerevisiae Hog1 (https://swissmodel.expasy.org/repository/uniprot/P32485?csm=D95A20242587CE06).

## Funding information

Work at the IR-T lab was supported by grants (BFU2017-90114-P and PID2020-120609GB-I00) by MCIN/AEI/ 10.13039/501100011033 and “ERDF A way of making Europe”. To JJS-E and AC, work was funded by the MCIN/AEI/10.13039/501100011033 (PID2019-108349RB-100). ED-M received a MEFP FPU fellowship FPU16/05237.

---

**Figure S1.**
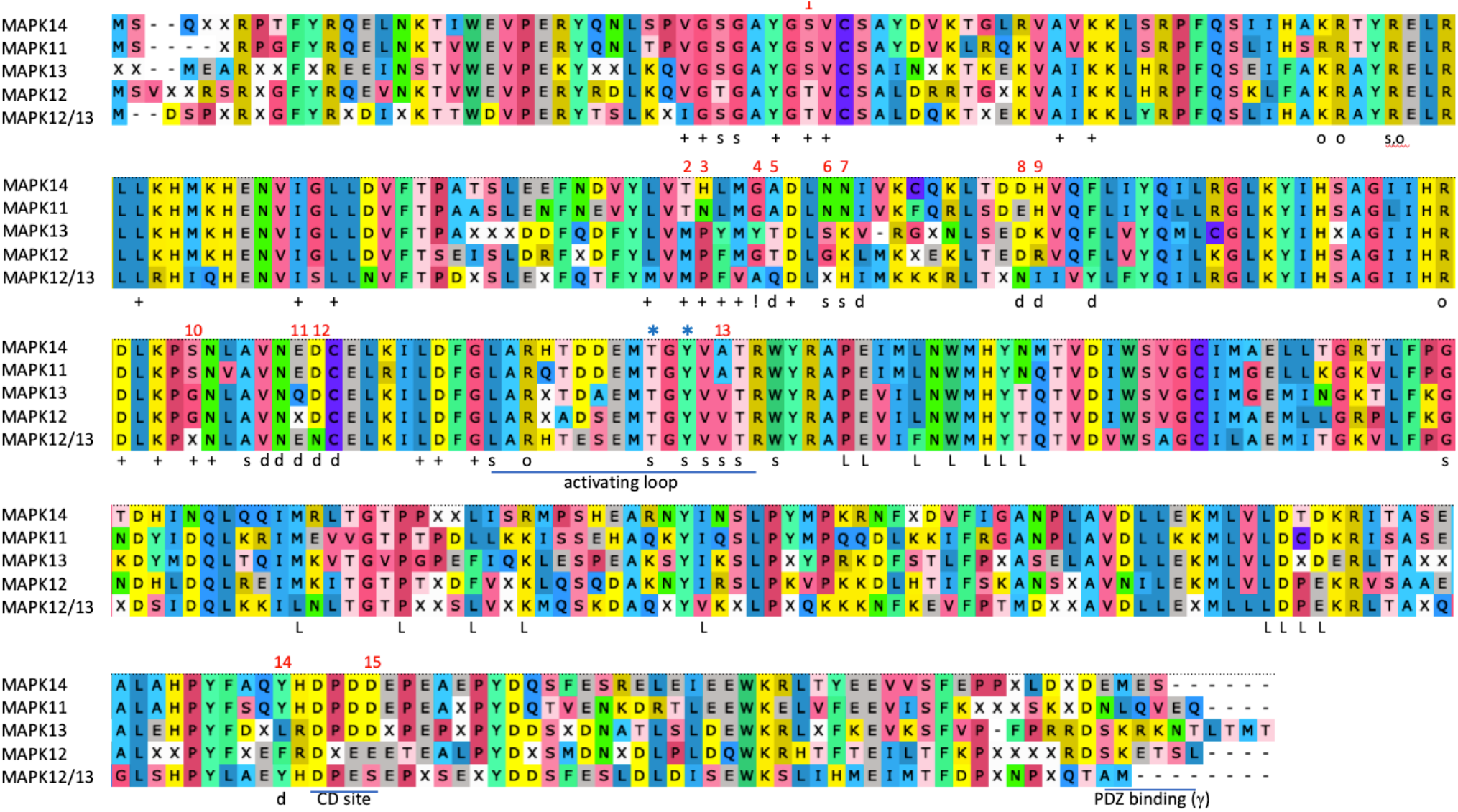
Alignment of the p38 consensus sequences of non-mammalian vertebrate lineages. Designation: (+) ATP binding site; (s) substrate binding site; (o) AA interacts with the conserved threonine when phosphorylated, necessary for folding into active state conformation; (d) docking site, (L) lipid binding site; (!) hinge point, the two domains rotate around hinge point during activation; asterisks mark the specific to the stress MAP kinases phosphorylation sites; PDZ binding domain is conserved only in p38γ. Red numbers point at the AA substitutions in functionally important sites.

## Notes

### Competing Interest Statement

The authors have declared no competing interest.

### Summary of Updates

Corrected the typo in NaCl concentrations. Added a supplementary Table S7 (cells viability assay in hyper-osmotic conditions)

